# DeepGOZero: Improving protein function prediction from sequence and zero-shot learning based on ontology axioms

**DOI:** 10.1101/2022.01.14.476325

**Authors:** Maxat Kulmanov, Robert Hoehndorf

## Abstract

**Motivation:** Protein functions are often described using the Gene Ontology (GO) which is an ontology consisting of over 50,000 classes and a large set of formal axioms. Predicting the functions of proteins is one of the key challenges in computational biology and a variety of machine learning methods have been developed for this purpose. However, these methods usually require significant amount of training data and cannot make predictions for GO classes which have only few or no experimental annotations.

**Results:** We developed DeepGOZero, a machine learning model which improves predictions for functions with no or only a small number of annotations. To achieve this goal, we rely on a model-theoretic approach for learning ontology embeddings and combine it with neural networks for protein function prediction. DeepGOZero can exploit formal axioms in the GO to make zero-shot predictions, i.e., predict protein functions even if not a single protein in the training phase was associated with that function. Furthermore, the zero-shot prediction method employed by DeepGOZero is generic and can be applied whenever associations with ontology classes need to be predicted.

**Availability:** http://github.com/bio-ontology-research-group/deepgozero

## 1 Introduction

Proteins are building blocks of any biological system, and to understand the biological system and its behavior on a molecular scale it is necessary to understand the functions of the proteins. Experimental identification of protein functions is time consuming and resource intensive. Next generation sequencing technologies have lead to a significant increase of the number of available DNA and protein sequences, thereby amplifying the challenge of identifying protein functions. Combined with the results of targeted experimental studies, computational researchers developed methods which analyze and predict protein structures (Baek *et al*., 2021; Jumper *et al*., 2021), determine the interactions between proteins (Pan *et al*., 2021; Sledzieski *et al*., 2021) and predict protein functions (Radivojac *et al*., 2013) based on the protein amino acid sequences. Several deep learning approaches were developed and applied to protein function prediction (Kulmanov *et al*., 2017; Kulmanov and Hoehndorf, 2019; Cao and Shen, 2021; You *et al*., 2021); however there are still several challenging questions that remain unanswered in computationally predicting protein functions.

For function prediction methods, there are two key challenging questions. First, it remains challenging to determine how a computational model can learn efficiently from protein sequences and effectively combine protein sequences with other sources of information such as the protein structure, interactions, or literature. Second, it remains challenging to predict the correct set of functions in the large, complex, unbalanced and hierarchical space of biological functions. Functions are described using the Gene Ontology (GO) (Ashburner *et al*., 2000) which is a large ontology with over 50,000 classes. It contains three sub-ontologies: the Molecular Functions Ontology (MFO), the Biological Processes Ontology (BPO) and the Cellular Components Ontology (CCO). The classes in these three ontologies are not independent and stand in formally defined relations that need to be considered while making predictions.

Many GO classes have never been used to characterize the functions of a protein, or only few proteins have been characterized with a particular function. The low number of annotations (or the absence of annotations) makes it challenging to directly train a machine learning model to predict these functions, or to make similarity-based predictions for these classes. More than 20, 000 classes in the GO have fewer than 100 proteins annotated (based on experimental evidence), and these classes are often specific and therefore highly informative classes. However, many of those classes have been formally defined by reusing other classes and relations using Description Logic axioms (Baader *et al*., 2003). For example, *regulation of binding* (`GO:0051098`) is defined as *biological regulation* (`GO:0065007`) and *regulates* (`RO:0002211`) some *binding* (`GO:0005488`). Using this definition we can annotate to the *regulation of binding* class if we know that proteins contribute to a *biological regulation* that regulates *binding*. GO has around 12, 000 definition axioms and more than 500, 000 other axioms which can be utilized to make predictions for classes without any, or with only few (less than 100), experimental annotations.

We developed DeepGOZero which combines a model-theoretic approach for learning ontology embeddings and protein function prediction. The background knowledge in GO in the form of formal axioms helps DeepGOZero to improve function prediction performance for specific classes (i.e., classes with few annotated proteins) and enables zero-shot predictions for classes without any annotations.

We evaluate the performance of DeepGOZero using both protein-centric and class-centric metrics (Radivojac and Clark, 2013); protein-centric metrics determine how accurate and complete the predictions are for proteins, and the class-centric metrics determine how reliable a particular function can be predicted. We demonstrate that DeepGOZero can achieve comparable performance to the state of the art in protein-centric metrics and outperforms all our baseline methods in the class-centric evaluations. We show that DeepGOZero performs best for classes with very few annotations and results in a strong predictive signal in zero-shot predictions. We compare DeepGOZero with state of the art methods such as NetGO2.0, DeepGraphGO and TALE+, all of which combine different types of information about proteins. We find that DeepGOZero can achieve comparable or better performance even based on only sequence based features. DeepGOZero is freely available at http://github.com/bio-ontology-research-group/deepgozero.

## 2 Methods

### 2.1 Materials and Data

In our experiments, we use two datasets that are generated using two different strategies. First, we downloaded the UniProt/SwissProt Knowledgebase (UniProtKB-SwissProt) (The UniProt Consortium, 2018) version 2021_04 released on 29-Sep-2021. We filtered all proteins with experimental functional annotations with evidence codes EXP, IDA, IPI, IMP, IGI, IEP, TAS, IC, HTP, HDA, HMP, HGI, HEP. The dataset contains 77, 647 reviewed and manually annotated proteins. For this dataset we use Gene Ontology (GO) released on 2021-11-16. We train and evaluate models for each of the sub-ontologies of GO separately.

The UniProtKB-SwissProt dataset includes proteins from over two thousands different organisms. Many proteins are orthologous and perform similar functions. For example, proteins `1433E_HUMAN`, `1433E_MOUSE` and `1433E_RAT` are identical in their sequence and share almost the same functional annotations. We computed sequence similarity using Diamond (Buchfink *et al*., 2014) and found that there are over 70,000 pairs of different proteins with more than 80% sequence identity. This means that, on average, every protein has a very similar other protein in the dataset. Figure 1 plots the similarity of all pairs of proteins.

**Fig. 1.**
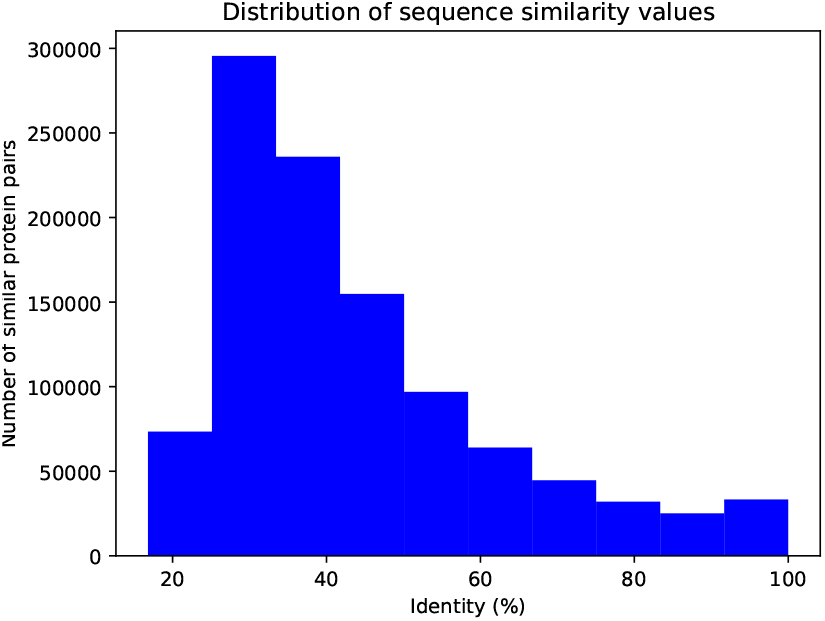
Distribution of proteins by sequence similarity. We compute pairwise similarity between all proteins in the SwissProt dataset and the figure shows the frequency of pairs by sequence similarity.

To train our prediction models on this dataset and ensure that it generalizes well to novel proteins, we split the proteins into training, validation and testing sets. If we split proteins randomly, the generated splits will contain proteins that are very similar or almost identical. Using such datasets in machine learning may lead to an overfitting problem and the prediction models will not generalize well (Tetko *et al*., 1995). Therefore, we decided to group the proteins by their similarity before generating a random split. We place proteins with sequence identity over 50% into the same groups and used 81% of the groups for training, 9% for validation and 10% for testing. Table 1 provides a summary of the dataset.

**Table 1.** Summary of the UniProtKB-SwissProt dataset

| Ontology | Terms | Proteins | Groups | Training | Validation | Testing |
| --- | --- | --- | --- | --- | --- | --- |
| MFO | 6,868 | 43,279 | 22,445 | 34,716 | 3,851 | 4,712 |
| BPO | 21,381 | 58,729 | 29,311 | 47,733 | 5,552 | 5,444 |
| CCO | 2,832 | 59,257 | 30,006 | 48,318 | 4,970 | 5,969 |

The table shows the number of GO terms, total number of proteins, number of groups of similar proteins, number of proteins in training, validation and testing sets for the UniProtKB-SwissProt dataset.

To compare our method with the state-of-art function prediction methods such as NetGO2.0 (Yao *et al*., 2021), DeepGraphGO (You *et al*., 2021) and TALE+ (Cao and Shen, 2021) we use the dataset generated by the authors of the NetGO2.0 method. The NetGO2.0 authors generate the dataset splits based on the time of annotation following the CAFA (Radivojac *et al*., 2013) challenge rules. They downloaded experimentally annotated proteins from GOA (The Gene Ontology Consortium, 2018) and UniProtKB including unreviewed proteins and generated training, validation and testing splits based on the following dates:

- Training – all data annotated in December 2018 or before
- Validation – proteins experimentally annotated from January 2019 to January 2020 and not before January 2019
- Testing – proteins experimentally annotated between February 2020 and October 2020 and not before February 2020

Experimental annotations were filtered based on evidence codes `EXP`, `IDA`, `IPI`, `IMP`, `IGI`, `IEP`, `TAS`, `IC` and the testing set proteins include only 17 species from the CAFA4 challenge targets. In addition, proteins which have been annotated with only the “protein binding” (`GO:0005515`) class were removed from the testing set to unbias it according to CAFA challenge instructions. This dataset does not consider sequence similarity. Table 2 summarizes the dataset.

**Table 2.** Summary of the NetGO dataset

| Ontology | Terms | Training | Validation | Testing |
| --- | --- | --- | --- | --- |
| MFO | 6,854 | 62,646 | 1,128 | 505 |
| BPO | 21,814 | 89,828 | 1,124 | 491 |
| CCO | 2,880 | 81,377 | 1,359 | 268 |

The table shows the number of GO terms and the number of proteins used as part of training, validation, and testing in the NetGO dataset.

### 2.2 Baseline methods

#### 2.2.1 DiamondScore

The DiamondScore method is based on the sequence similarity score obtained by Diamond (Buchfink *et al*., 2014). The method aims to find similar sequences from the training set and transfer their annotations. We use the normalized bitscore to compute the prediction score for a query sequence *q*:

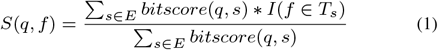

where *E* is a set of similar sequences filtered by e-value of 0.001, *T*_*s*_ is a set of true annotations of a protein with sequence *s* and *I* is an indicator function which returns 1 if the condition is true and 0 otherwise.

#### 2.2.2 MLP

The MLP method predicts protein functions using multi-layer perceptron (MLP) networks from a protein sequence’s InterPro domain annotations obtained with InterProScan (Mitchell *et al*., 2014). We represent a protein with a binary vector for all the InterPro domains and pass it to two layers of MLP blocks where the output of the second MLP block has residual connection to the first block. This representation is passed to the final classification layer with sigmoid activation function. One MLP block performs the following operations:

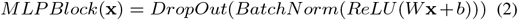

The input vector x of length 26, 406, which represents InterPro domain annotations, is reduced to 1, 024 by the first MLPBLock:

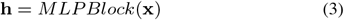

Then, this representation is passed to the second MLPBlock with the input and output size of 1, 024 and added to itself using residual connection:

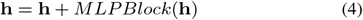

Finally, we pass this vector to a classification layer with a sigmoid activation function. The output size of this layer is the same as the number of classes in each subontology:

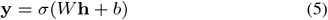

We train a different model for each subontology in GO.

#### 2.2.3 DeepGOPlus and DeepGOCNN

DeepGOPlus (Kulmanov and Hoehndorf, 2019) predicts functional annotations of proteins by combining DeepGOCNN, which predicts functions from the amino acid sequence of a protein using a 1-dimensional convolutional neural network, with the DiamondScore method. DeepGOCNN captures sequence motifs that are related to GO functions.

### 2.3 NetGO2.0

NetGO2.0 (Yao *et al*., 2021) is an ensemble method which combines five different sources of information. Specifically, NetGO2.0 uses the GO term frequency, sequence features (InterPro domains), protein–protein interaction networks, a recurrent neural network (RNN), and literature. This method is an improved version of the GOLabeler (You *et al*., 2018) method which was the best-performing method in the CAFA3 challenge. We used the NetGO2.0 webserver to obtain predictions for the test sets.

### 2.4 DeepGraphGO

The DeepGraphGO (You *et al*., 2021) method uses a neural network to combine sequence features with PPI networks by using graph convolutional neural networks. We obtained predictions for our test sets by using the source code provided by the authors of this method.

### 2.5 TALE+

TALE+ (Cao and Shen, 2021) predicts functions using a transformer-based deep neural network model. TALE+ also uses a hierarchical loss function to encode the taxonomy of the GO into the model. The deep neural network predictions are combined with predictions based on sequence similarity, similarly to DeepGOPlus.

### 2.6 DeepGOZero

We call our model DeepGOZero because it allows us to predict functional annotations for GO classes that do not have any training proteins. To achieve this goal, we combine protein function prediction with a model-theoretic approach for embedding ontologies into a distributed space, ELEmbeddings (Kulmanov *et al*., 2019). We use InterPro domain annotations represented as binary vector as input and apply two layers of MLPBlock from our MLP baseline method to generate an embedding of size 1024 for a protein. For a given protein *p* we predict annotations for a class *c* using the following formula:

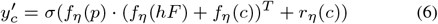

where *f*_*η*_ is an embedding function, *hF* is the `hasFunction` relation, *r*_*η*_ (*c*) is the radius of an *n*-ball for a class *c* and *σ* is a sigmoid activation function.

We compute the binary crossentropy loss between our predictions and the labels, and optimize them together with four normal form losses for ontology axioms from ELEmbeddings. We use the Adam (Kingma and Ba, 2014) optimizer to minimize the following loss function:

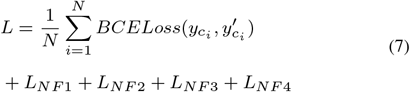

ELEmbeddings use normalized axioms in the following four different forms:

- *NF* 1 : *C* ⊑ *D*
- *NF* 2 : *C* ⊓ *D* ⊑ *E*
- *NF* 3 : *C* ⊑ ∃*R*.*D*
- *NF* 4 : ∃*R*.*C* ⊑ *D*

where *C, D, E* represent classes and *R* represents relations in the ontology. We convert the GO axioms into these four normal forms using a set of conversion rules implemented in the JCel reasoner (Mendez, 2012). As a result, we obtain a set of axioms all of which are in one of these four normal forms, and which are equivalent to the axioms in GO. ELEmbeddings uses these normalized GO axioms as constraints and projects each GO class into an *n*-ball (represented as a center point in *n*-dimensional space and a radius) and each relation as a transformation within *n*-dimensional space. We set the dimension *n* to 1024 so that it is the same as our protein embeddings. We compute the losses for each of the normal forms using the following formulas:

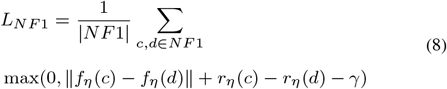

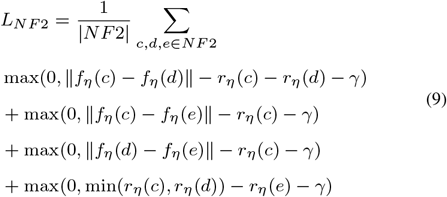

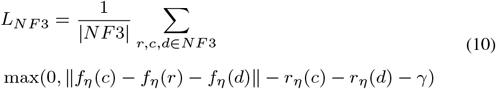

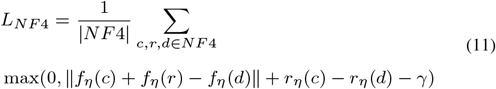

The parameter *γ* is a margin parameter and ‖ ‖ is the *L*_2_ norm of the vectors. In addition, we use batch normalization for class embedding vectors to regularize the model and avoid overfitting.

### 2.7 Evaluation

To evaluate our predictions we use the CAFA (Radivojac *et al*., 2013) protein-centric evaluation metrics *F*_max_ and *S*_min_ (Radivojac and Clark, 2013). In addition, we report the area under the precision-recall curve (AUPR) and class-centric average area under the receiver operating characteristic curve (AUROC) which are reasonable measures for evaluating predictions with high class imbalance (Davis and Goadrich, 2006).

*F*_max_ is a maximum protein-centric F-measure computed over all prediction thresholds. First, we compute average precision and recall using the following formulas:

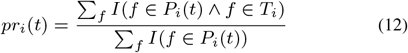

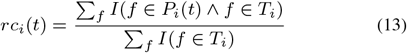

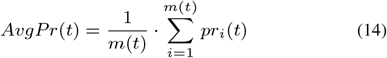

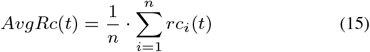

where *f* is a GO class, *T*_*i*_ is a set of true annotations, *P*_*i*_(*t*) is a set of predicted annotations for a protein *i* and threshold *t, m*(*t*) is a number of proteins for which we predict at least one class, *n* is a total number of proteins and *I* is an indicator function which returns 1 if the condition is true and 0 otherwise. Then, we compute the *F*_max_ for prediction thresholds *t* ∈ [0, 1] with a step size of 0.01. We count a class as a prediction if its prediction score is greater or equal than *t*:

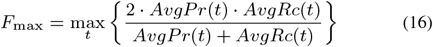

*S*_min_ computes the semantic distance between real and predicted annotations based on information content of the classes. The information content *IC*(*c*) is computed based on the annotation probability of the class *c*:

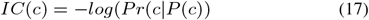

where *P* (*c*) is a set of parent classes of the class *c*. The *S*_min_ is computed using the following formulas:

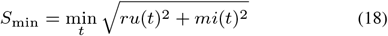

where *ru*(*t*) is the average remaining uncertainty and *mi*(*t*) is average misinformation:

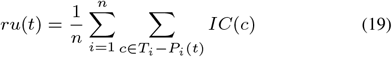

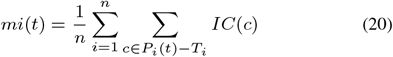

## 3 Results

### 3.1 DeepGOZero

The aim of DeepGOZero is to use the background knowledge contained in the Description Logic axioms of GO to improve protein function prediction. We hypothesize that ontology axioms will help to improve the quality of predictions and allow us to predict functional annotations for ontology terms without training samples (zero-shot) using only the ontology axioms, thereby combining neural and symbolic AI methods within a single model (Mira *et al*., 2003). Axioms in the GO formally constrain classes (The Gene Ontology Consortium, 2018). The simplest form of axiom are subclass axioms, such as *Ion binding* (`GO:0043167`) SubClassOf *Binding* (`GO:0005488`), which indicates that every instance of *Ion binding* must also be an instance of *Binding* (Smith *et al*., 2005). These axioms also constrain annotations of proteins; if a protein *P* has function *Ion binding*, it will also have the function *Binding* (in virtue of all *Ion binding* instances being *Binding* instances). In the context of GO, this rule is also known as the “true path rule” (Ashburner *et al*., 2000) and this rule is exploited by several protein function prediction methods (Radivojac *et al*., 2013); in most cases, exploiting the true path rule for function prediction can improve prediction performance (Radivojac *et al*., 2013; Kulmanov *et al*., 2017).

However, there are axioms beyond subclass axioms in the GO (The Gene Ontology Consortium, 2018). For example, the class *serine-type endopeptidase inhibitor activity* (`GO:0004867`) is defined as being equivalent to *molecular function regulator* (`GO:0098772`) that *negatively regulates* (`RO:0002212`) some *serine-type endopeptidase activity* (`GO:0004252`). Based on this axiom, to predict whether a protein *P* has the function *serine-type endopeptidase inhibitor activity*, it suffices to know that the protein is a *molecular function regulator* and can negatively regulate *serine-type endopeptidase activity*. This axiom can therefore be used in two ways by a function prediction model. First, it imposes an additional constraint on functions predicted for *P*, and this constraint may both reduce search space during optimization and improve prediction accuracy. Second, the axiom can allow us to predict the function *serine-type endopeptidase inhibitor activity* for a protein *P* even if not a single protein seen during training has this function. To achieve this zero-shot prediction goal, the model must be able to predict that a protein *P* has the function *molecular function regulator* and the *negatively regulates* relation of the protein *P* to *serine-type endopeptidase activity*, and be able to combine both into the compound prediction *serine-type endopeptidase inhibitor activity*. For zero-shot predictions, we recognize that both *molecular function regulator* and *serine-type endopeptidase activity* are “simpler” classes in the sense that they are more general and therefore have more annotations than the compound class (in virtue of the “true path rule”); and, similarly, the relation *negatively regulates* is used across multiple GO classes and has therefore more associated proteins than the compound class.

We designed DeepGOZero to predict functions for proteins using the GO axioms. Specifically, we first use the geometric ontology embedding method EL Embeddings (Kulmanov *et al*., 2019) to generate a space in which GO classes are *n*-balls in an *n*-dimensional space and the location and size of the *n*-balls are constrained by the GO axioms. The ontology axioms are only used during the training phase of DeepGOZero to generate the space constrained by GO axioms (see Section 2.6). We then use a neural network to project proteins into the same space in which we embedded the GO classes and predict functions for proteins by their proximity and relation to GO classes. Specifically, DeepGOZero uses as input the InterPro domain annotations of a protein where we represent the annotations as a binary vector. The binary vector of InterPro domain annotations are processed by MLP layers to generate an embedding vector of the same size of an embedding vector for GO classes. Then, DeepGOZero jointly minimizes prediction loss for protein functions and the ELEmbeddings loss that impose constraints on the classes (see Section 2.6). Figure 2 provides an example of the predictions made by DeepGOZero.

**Fig. 2.**
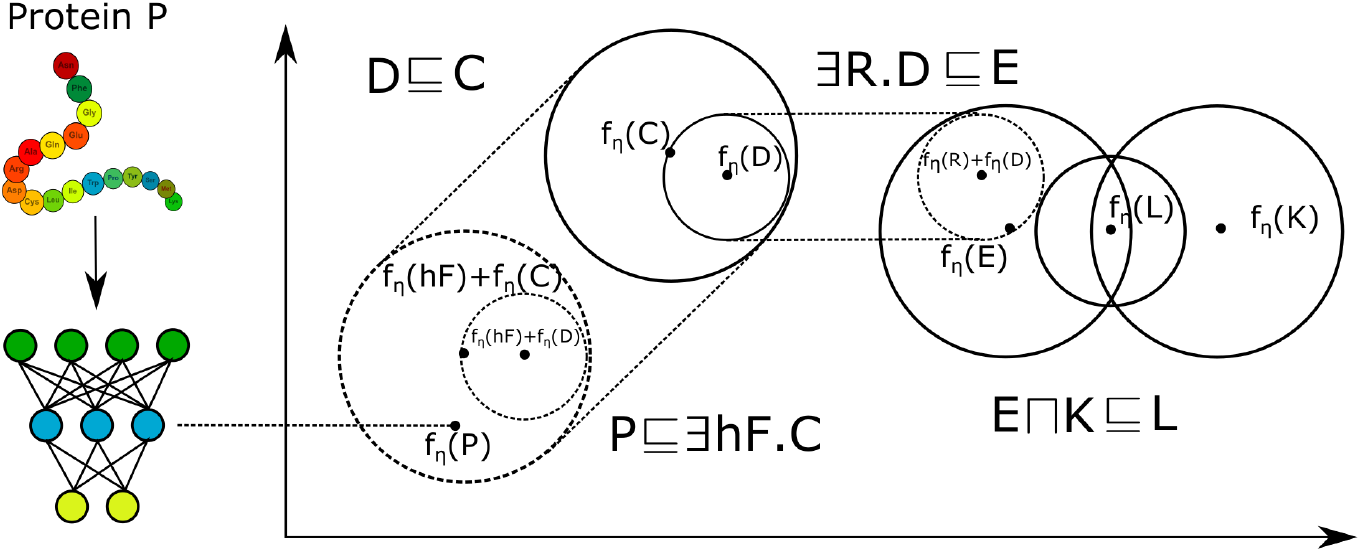
The figure provides a high-level overview and example of the DeepGOZero model. On the left, a protein *P* is embedded in a vector space using an MLP whereas the right side shows how GO axioms are embedded using the EL Embedding method; the MLP embeds the protein in the same space as the GO axioms. *C, D, E, K*, and *L* are classes defined by a center point and radius, *R* is a relation used in an axiom in GO, and the set of GO axioms constrain the geometric relations of the classes (for example, the class ∃*R*.*D* is the transformation of *D* by the relation *R, f*_*η*_ (∃*R*.*D*) = *f*_*η*_ (*R*) + *f*_*η*_ (*D*)). Function prediction for a protein is based on the hasFunction (*hF*) relation of the protein to the GO function embeddings. Both the protein and the GO class embeddings are optimized jointly during training of DeepGOZero.

We apply our model to the proteins in our UniProtKB-SwissProt dataset testing set and evaluate the predictive performance using the CAFA measures. We evaluate the performance in the three branches of GO separately as they have different characteristics in terms of number and type of axioms and also can benefit differently from the features (InterPro domains) we use in DeepGOZero. Table 3 shows the results for MFO, Table 4 for BPO, and Table 5 for CCO. We find that, in protein centric evaluations, MLP and DeepGOZero methods perform significantly better than other baseline methods when combined with DiamondScore in MFO and BPO evaluations in terms of *F*_max_ and *S*_min_ whereas DeepGOPlus scores the best *F*_max_ in CCO evaluations. MLP + DiamondScore is slightly better than DeepGOZero + DiamondScore in terms of *F*_max_ and *S*_min_, but DeepGOZero + DiamondScore has a better AUPR in MFO and CCO evaluations. Noticably, in the class-centric evaluation, DeepGOZero + DiamondScore has the best AUC in all evaluations.

**Table 3.**
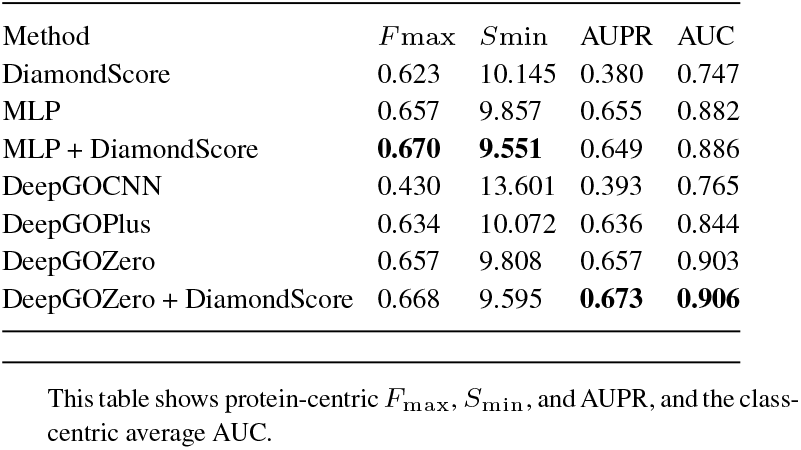
Prediction results for Molecular Function on the UniProtKB-SwissProt dataset

**Table 4.**
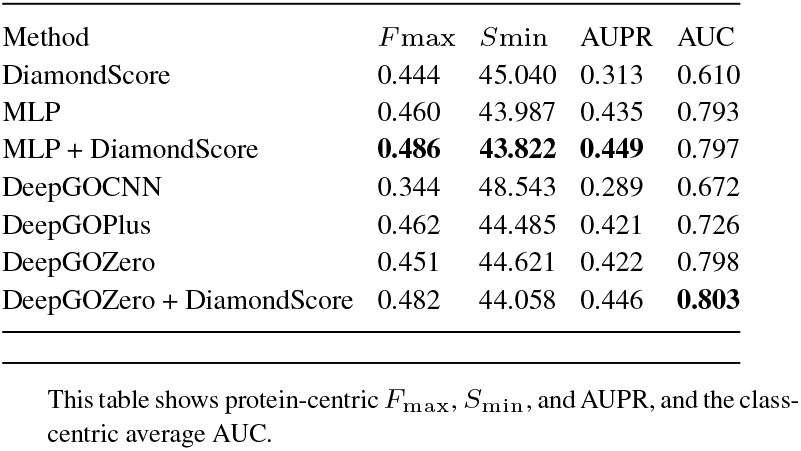
Prediction results for Biological Process on the UniProtKB-SwissProt dataset

**Table 5.** Prediction results for Cellular Component on the UniProtKB-SwissProt dataset

| Method | $F_{\max}$ | $S_{\min}$ | AUPR | AUC |
| --- | --- | --- | --- | --- |
| DiamondScore | 0.581 | 11.092 | 0.352 | 0.648 |
| MLP | 0.667 | **10.523** | 0.670 | 0.846 |
| MLP + DiamondScore | 0.666 | 10.526 | 0.654 | 0.851 |
| DeepGOCNN | 0.641 | 11.396 | 0.645 | 0.775 |
| DeepGOPlus | **0.672** | 10.591 | 0.667 | 0.821 |
| DeepGOZero | 0.661 | 10.681 | 0.665 | 0.854 |
| DeepGOZero + DiamondScore | 0.667 | 10.615 | **0.701** | **0.860** |

This table shows protein-centric $F_{\max}$ , $S_{\min}$ , and AUPR, and the class-centric average AUC.

We further analyze term centric prediction performance of the MLP and DeepGOZero methods based on the specificity of the GO class which we define by the number of their annotations. We compute the average AUC of testing set predictions and group them by the number of all annotations for each term. We find that DeepGOZero, on average, has better predictive performance for GO classes with less than 50 annotations whereas the MLP performs better for GO classes that have more annotations. Figure 3 shows this comparison of the DeepGOZero and MLP methods.

**Fig. 3.**
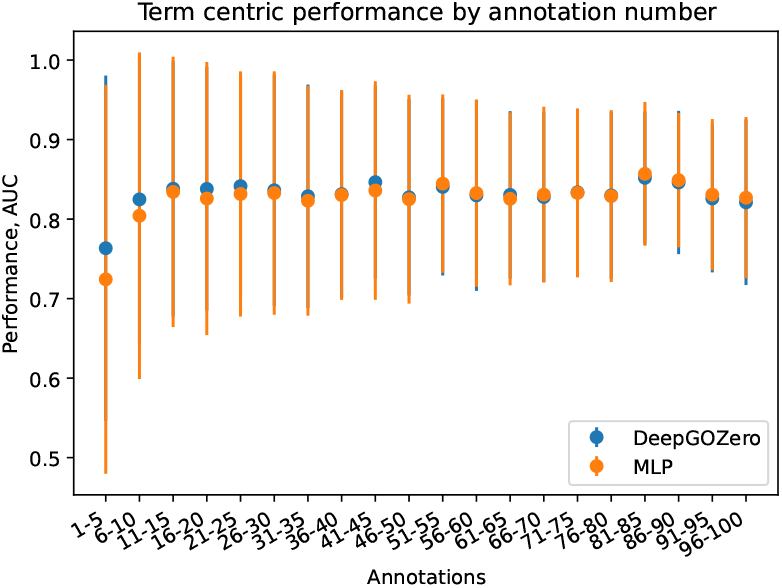
Average prediction performance of classes grouped by annotation size on UniprotKB-Swissprot dataset

We compare our method with the state of the art function prediction methods NetGO2.0 (Yao *et al*., 2021), DeepGraphGO (You *et al*., 2021) and TALE+ (Cao and Shen, 2021); for this purpose, we train our models on the dataset generated by NetGO2.0 authors. This dataset was generated by following the CAFA (Radivojac *et al*., 2013) rules, and DeepGraphGO and TALE+ methods were not trained on the testing set of this dataset. We obtain predictions of DeepGraphGO and TALE+ using their source codes and use the webserver of NetGO2.0 as the source code is not available. We evaluate the predictions using our evaluation measures.

We find that in the MFO evaluation, DeepGOZero + DiamondScore has the best class-centric average AUC and NetGO2.0 performs best in all protein-centric evaluations. For BPO, our MLP baseline combined with DiamondScore achieves the best protein-centric *F*_max_ and AUPR, whereas in class-centric evaluation DeepGraphGO performs the best. Also, DeepGraphGO provides best predictions in terms of protein-centric *F*_max_ and *S*_min_ in the CCO evaluation. Finally, TALE+ achieves the best AUPR and DeepGOPlus achieves the best class-centric average AUC in CCO evaluation. Table 6 summarizes the results for this experiment. Overall, the evaluation shows that DeepGOZero can achieve a predictive performance comparable to the state of the art function prediction methods although DeepGOZero uses only sequence-based features.

**Table 6.**
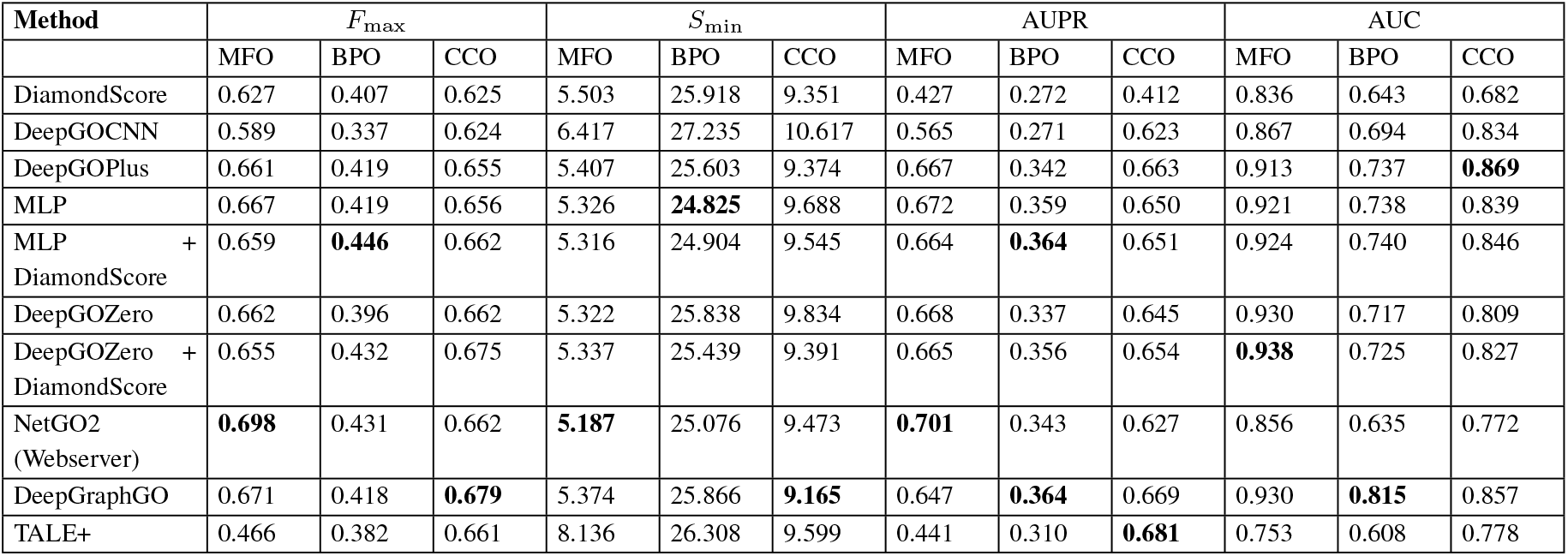
The comparison of performance on the NetGO dataset.

We further compare the different methods in terms of the specificity of their best predictions and find that DeepGOZero+DiamondScore provides the predictions with the highest average information content in BPO and CCO evaluations. In the MFO evaluation, TALE+ provides the most specific annotations (see Table 7). While the BPO and CCO branches of GO have a large number of formal definitions and axioms, the MFO branch has only a low number of these definitions, and the lower specificity of predicted classes for DeepGOZero in MFO indicates that the axioms are important to improve specificity of the predictions.

**Table 7.**
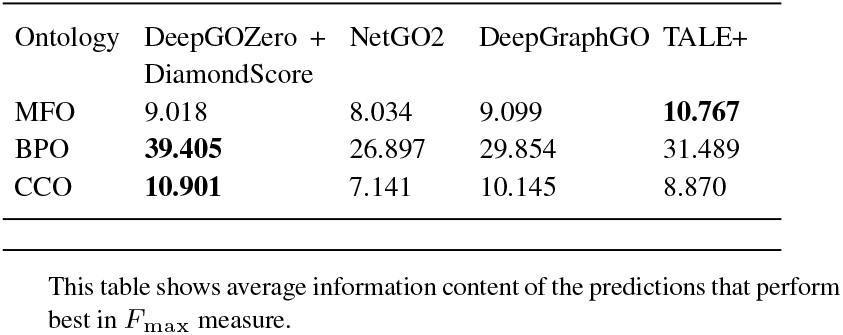
Average information content of the predictions on NetGO dataset

| Ontology | DeepGOZero + DiamondScore | NetGO2 | DeepGraphGO | TALE+ |
| --- | --- | --- | --- | --- |
| MFO | 9.018 | 8.034 | 9.099 | **10.767** |
| BPO | **39.405** | 26.897 | 29.854 | 31.489 |
| CCO | **10.901** | 7.141 | 10.145 | 8.870 |

This table shows average information content of the predictions that perform best in $F_{\max}$ measure.

### 3.2 Zero-shot function prediction

DeepGOZero makes predictions for protein functions based on geometric relations to embeddings in a vector space. As we have embeddings for all classes in the GO, DeepGOZero can predict functions for a protein even if not a single protein in the training set was associated with the function. We refer to these predictions as “zero-shot” function predictions.

To evaluate the performance of zero-shot predictions we perform two experiments on our similarity based splitted UniProtKB-SwissProt dataset. First, we test the performance of predictions for specific classes that have equivalent class axiom definitions. For example, the class *serine-type endopeptidase inhibitor activity* (`GO:0004867`) in the GO is defined as being equivalent to *molecular function regulator* (`GO:0098772`) and *negatively regulates* (`RO:0002212`) some *serine-type endopeptidase activity* (`GO:0004252`). For our first experiment, we use only the classes which have 100 or more annotations in order to compare zero-shot prediction performance with the performance of the trained model where we use all GO classes that have experimental protein annotations. We find 16 such classes and remove their annotations from our UniProtKB-SwissProt dataset (before propagating annotations using the true path rule). We train new models of DeepGOZero for each subontology after removing these annotations and predict using the embeddings only (i.e., DeepGOZero has never seen a protein annotated to these classes during training). We perform two different tests: first, we evaluate the performance on proteins which have never been seen during training (test set performance); and, second, because we remove the GO class from the annotations of all proteins (training and test set), we test whether and how well DeepGOZero can predict this class for all proteins. We compare the performance directly with the performance of the supervised model where the class was used during model training. Table 8 summarizes the performance of zero-shot predictions for these specific classes (using the class-centric evaluation measures). We find that, while the zero-shot predictions are substantially better than random, training consistently improves over the zero-shot prediction performance. However, zero-shot predictions achieve an average AUC of 0.745, demonstrating a strong predictive signal despite being lower than performance in supervised predictions. Furthermore, we find that zero-shot predictions after training on the proteins (but not the class) can also improve performance compared to zero-shot predictions where the protein also has not been included in training. DeepGOZero embeds proteins in the ELEmbedding space, and the improved performance when training on the protein shows that the protein is placed closer to its “correct” position within that space based on other annotations compared to the first test where no information about the protein is available at all.

**Table 8.** Zero-shot and trained prediction performance on specific classes with more than 100 annotations. Evaluation measures are class-centric. AUC(test) is the zero-shot performance on the test set, i.e., neither the class nor the protein were included during model training; AUC(all) is the zero-shot performance on all proteins, i.e., the class was never seen during training but the model has seen the proteins (annotated with other classes) during training; and AUC(trained) is the performance of the model on the testing set when trained with the class (i.e., the protein is not seen but other proteins with the class were used during training).

| Ontology | Term | Name | AUC (test) | AUC (all) | AUC (trained) |
| --- | --- | --- | --- | --- | --- |
| MFO | GO:0001227 | DNA-binding transcription repressor activity, RNA polymerase II-specific | 0.257 | 0.405 | 0.932 |
| MFO | GO:0001228 | DNA-binding transcription activator activity, RNA polymerase II-specific | 0.574 | 0.699 | 0.948 |
| MFO | GO:0003735 | structural constituent of ribosome | 0.400 | 0.194 | 0.940 |
| MFO | GO:0004867 | serine-type endopeptidase inhibitor activity | 0.972 | 0.967 | 0.985 |
| MFO | GO:0005096 | GTPase activator activity | 0.847 | 0.870 | 0.938 |
| BPO | GO:0000381 | regulation of alternative mRNA splicing, via spliceosome | 0.855 | 0.865 | 0.907 |
| BPO | GO:0032729 | positive regulation of interferon-gamma production | 0.870 | 0.919 | 0.928 |
| BPO | GO:0032755 | positive regulation of interleukin-6 production | 0.719 | 0.819 | 0.885 |
| BPO | GO:0032760 | positive regulation of tumor necrosis factor production | 0.861 | 0.906 | 0.918 |
| BPO | GO:0046330 | positive regulation of JNK cascade | 0.855 | 0.894 | 0.894 |
| BPO | GO:0051897 | positive regulation of protein kinase B signaling | 0.772 | 0.864 | 0.890 |
| BPO | GO:0120162 | positive regulation of cold-induced thermogenesis | 0.637 | 0.789 | 0.730 |
| CCO | GO:0005762 | mitochondrial large ribosomal subunit | 0.889 | 0.975 | 0.874 |
| CCO | GO:0022625 | cytosolic large ribosomal subunit | 0.898 | 0.969 | 0.893 |
| CCO | GO:0042788 | polysomal ribosome | 0.858 | 0.950 | 0.889 |
| CCO | GO:1904813 | ficolin-1-rich granule lumen | 0.653 | 0.782 | 0.792 |
|  |  | Average | 0.745 | 0.804 | 0.896 |

Furthermore, we evaluate the performance of zero-shot predictions for classes with a very low number (*<* 10) of training samples; these classes will benefit the most from zero-shot predictions as they are rarely included in supervised training due to the low number of annotations. We first evaluate the average class-centric AUC for all classes and find that DeepGOZero achieves an AUC of 0.804, 0.737 and 0.819 for the MFO, BPO, and CCO subontologies. Then, we filter the classes that have definition axioms in GO and find that their predictions are significantly better than the performance for all classes with AUCs of 0.862, 0.786 and 0.915 for MFO, BPO, and CCO, respectively. Table 9 provides the number of classes and average AUCs for zero-shot predictions with DeepGOZero. In total, there are 13, 501 class without any annotations and 17, 375 classes with 58, 770 annotations. Figure 4 shows the distribution of classes and average class-centric AUC for zero-shot predictions by their number of annotations.

**Table 9.** Zero-shot prediction performance on classes with less than 10 annotations

| Ontology | All classes |  | Defined classes |  |
| --- | --- | --- | --- | --- |
|  | Num. classes | AUC | Num. classes | AUC |
| MFO | 4,791 | 0.804 | 95 | 0.862 |
| BPO | 11,092 | 0.737 | 4,598 | 0.786 |
| CCO | 1,492 | 0.819 | 151 | 0.915 |

The table shows average performance for all evaluated classes and classes that have definition axioms. It provides number of classes and average class-centric AUC.

**Fig. 4.**
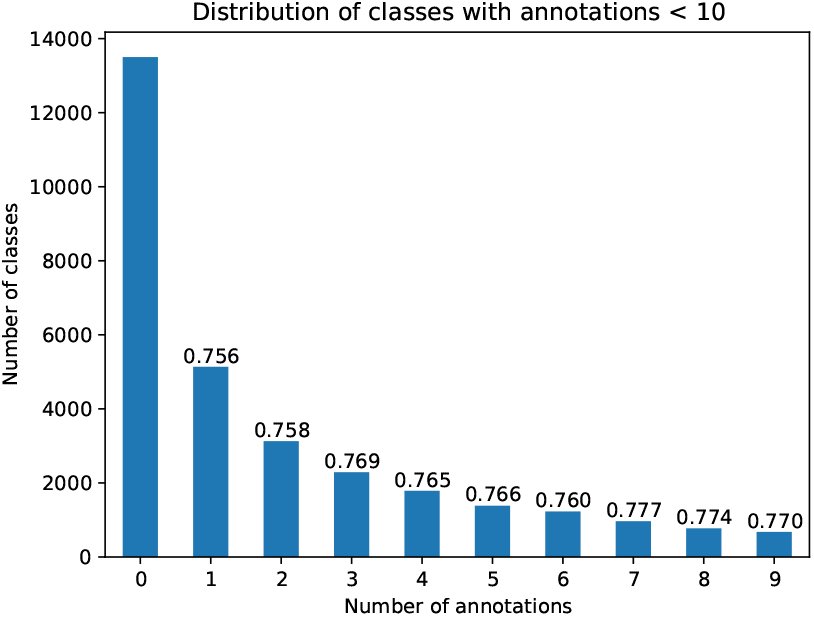
Distribution of GO classes by their number of annotated proteins (*<* 10) for zero-shot prediction evaluation.

## 4 Discussion

The life science community has spent significant resources on developing biomedical ontologies that formally represent the basic types of entities that are investigated across domains in life sciences (Smith *et al*., 2007; Jackson *et al*., 2021), as well as the relations in which they stand to each other (Smith *et al*., 2005). These ontologies are used in almost every biomedical database to ensure interoperability and data integration. However, the axioms in the ontologies that formally express the intended meaning of a class are still rarely exploited despite the large amount of information they contain. Only recently were methods developed that can make use of some of these axioms, in particular in the form of ontology embeddings that project the axioms into a space where they can be combined with other types of data (Kulmanov *et al*., 2020). To our knowledge, DeepGOZero is the first method that uses the axioms of the GO to constrain function prediction methods. With the use of axioms (beyond the subclass axioms between named classes which are used in hierarchical classifiers), our method can not only improve prediction performance for specific classes, but also predict functions for proteins in the absence of any training data for the functions.

Function prediction is a multi-class multi-label problem with a hierarchical dependencies of classes. Standard protein-centric evaluation metrics consider the whole hierarchy while measuring the performance of prediction methods. This makes them biased towards general ontology classes, and methods which perform well on the general and frequently annotated classes have a good performance (Radivojac and Clark, 2013). However, predicting general classes is less useful than predicting specific classes because specific classes provide more information about the functions of a protein and its role within an organism. We have shown that DeepGOZero, on average, improves the predictions of specific classes with few annotations and even can make predictions for classes without any annotations. Furthermore, the presence of formalized axioms improves DeepGOZero predictions substantially.

The main impact of zero-shot predictions is on predicting functions for which only few, or no, proteins have yet been identified, and which consequently are not part of protein function prediction benchmark and evaluation datasets. Our DeepGOZero model predicts 13, 501 distinct GO classes that have not a single protein associated based on experimental evidence; of these, 2, 935 classes are constrained by formal axioms for which DeepGOZero performs substantially better than for classes that are not constrained by an axiom. None of these functions can be predicted by other models, and they cannot be predicted based on sequence similarity either due to the absence of any annotated proteins. For the same reasons, we cannot computationally evaluate the accuracy of these predictions; however, we make our predictions available for validation in the future.

The CAFA challenge evaluates and selects the top methods based on *F*_max_ measure (Radivojac and Clark, 2013). This metric is a kind of semantic similarity measure which takes the hierarchical structure of the ontology into account. However, this metric does not consider the specificity of the classes and gives all classes the same weight. This makes the *F*_max_ biased towards general classes with more annotated proteins. For example, simple frequency based predictors result in overall performance substantially better than random when measured using *F*_max_ (Radivojac *et al*., 2013; Jiang *et al*., 2016; Zhou *et al*., 2019). We believe that this can be solved by weighting the classes by their specificity using a class specificity measure. A weighted measure, such as the weighted *F*_max_ (Radivojac and Clark, 2013), would give more weight to specific classes; however, it is not currently used in the evaluations and consequently models do not optimize for it. In addition, CAFA reports *S*_min_ measure which is based on the information content. This metric is also biased towards general classes because general classes have small information content and minimum *S*_min_ is considered best. The methods which make specific predictions have, on average, more false positives in specific classes and result in higher *S*_min_. In the future, measures that do not only evaluate overall predictive performance but also the specificity (and therefore the utility) of the predictions should be employed alongside the current evaluation measures.

## 5 Conclusion

DeepGOZero predicts protein functions from amino acid sequence features and incorporates the background knowledge from the GO in the prediction process. We embed GO into a vector space using a geometric embedding model and project proteins into the same space using a neural network. GO axioms constrain the function prediction and the use of axioms improves prediction performance for specific GO classes with few annotations and allows us to perform zero-shot function prediction. Furthermore, we demonstrate that DeepGOZero can provide comparable performance with state of the art function predictors by only using a sequence features. We believe that the performance can be further improved by combining other sources of information such as protein structure, PPIs and literature. The ontology-based prediction method of DeepGOZero is generic and can be adapted to other ontologies and other applications as well.

## Data availability

All data underlying this work, including source code, is freely available at https://github.com/bio-ontology-research-group/deepgozero.

